# Effects of the expression of random sequence clones on growth and transcriptome regulation in *Escherichia coli*

**DOI:** 10.1101/2021.11.22.469564

**Authors:** Devika Bhave, Diethard Tautz

## Abstract

Comparative genomic analyses have provided evidence that new genetic functions can emerge out of random nucleotide sequences. Here, we apply a direct experimental approach to study the effects of plasmids harbouring random sequence inserts under the control of an inducible promoter. Based on data from previously described experiments dealing with the growth of clones within whole libraries, we extracted specific clones that had shown either negative or positive effects on relative cell growth. We analysed these individually with respect to growth characteristics and impact on the transcriptome. We find that candidate clones for negative peptides lead to growth arrest by eliciting a general stress response. Overexpression of positive clones, on the other hand, does not change the exponential growth rates of hosts, but they show a growth advantage over a neutral candidate clone when tested in direct competition experiments. Transcriptomic changes in positive clones are relatively moderate and specific to each clone. We conclude from our experiments that random sequence peptides are indeed a suitable source for the *de novo* evolution of genetic functions.

## 1. Introduction

The origin of novelty is a fundamental theme in evolutionary genetics. While the focus is often on changes in existing genes and their effects on the downstream pathways, comparative genomic analysis has shown that *de novo* evolution of new genes also contributes much to evolutionary innovations (reviewed in (Ding, Zhou, & Wang, 2012; Kaessmann, 2010; McLysaght & Guerzoni, 2015; Tautz & Domazet-Loso, 2011)). Different mechanisms are responsible for generating new genes, including segmental duplications, retrogene insertions, and de novo evolution from non-coding DNA. While the latter mechanism has initially been considered to be very unlikely (see review in (Tautz, 2014)), it has by now been documented for several well-studied cases (see (Bornberg-Bauer, Hlouchova, & Lange, 2021; Ruiz-Orera, Villanueva-Cañas, & Albà, 2020; Van Oss & Carvunis, 2019) for recent reviews). Nonetheless, how frequently such an emergence would occur remains a matter of active debate. A key question in this debate is: what fraction of random sequence open reading frames (ORFs) have the potential to interact with intracellular components and could consequently be subjected to positive selection to establish a novel function for an organism?

We have previously addressed this question by expressing libraries with random sequence ORFs in *Escherichia coli* and studying frequency changes of each clone while allowing growth under competitive conditions for several generations (Neme, Amador, Yildirim, McConnell, & Tautz, 2017). We found that about 70% of the total clones showed a consistent positive or negative effect on the growth across multiple experimental replicates. However, the results of this experiment were challenged due to the design based on an expression vector, which could have interfered with the interpretation of the frequency changes of clones due to a possible vector-specific negative effect on the host (Knopp & Andersson, 2018; Weisman & Eddy, 2017). It was argued that the empty vector drives a strong expression after induction with isopropyl β-d-1-thiogalactopyranoside (IPTG), which could be generally harmful to the host, or the small 38 amino-acid long peptides encoded by the multiple cloning site (MCS) region could be detrimental. By inserting random sequences into the MCS, it is possible to either underestimate the generally harmful effects of overexpression or, more specifically, the potentially harmful expression of the peptides (Knopp & Andersson, 2018). This would only impact the clones that showed a positive growth effect since they can be interpreted to alleviate the negative effect of the empty vector non-specifically. However, the overall effect sizes of several of these clones would argue against such an interpretation (Tautz & Neme, 2018).

Based on a re-examination of the data using a new analysis pipeline that included not only the fulllength peptides as in (Neme et al., 2017) but also all shorter versions of the peptides that were in the library, we found no evidence of a strong vector effect in the experiments (Fajardo & Tautz, 2021). The study also confirmed that approximately 16% of peptides positively impact the relative growth of the cells. Interestingly, this study showed that shorter sequences (between 8-20 amino acids) constitute the highest fraction of positive peptides after four growth cycles. However, given that these experiments were done in the context of the growth of the entire library of clones, clonal interference is expected to occur that can significantly affect the growth trajectories of individual clones (Gerrish & Lenski, 1998; Van den Bergh, Swings, Fauvart, & Michiels, 2018).

Therefore, a key for a better understanding of the effects of random sequence ORF expression in *E. coli* is to trace specific effects for individual clones. Here we study a set of such candidate ORFs derived from the original library. We test individual growth phenotypes and monitor transcriptome changes after induction of expression to get insights into their effects on the cells.

We find that the expression of the negative clones induces a relatively generic stress response in the host. In contrast, the positive clones show no stress response but clone-specific effects. We conclude that the random sequence ORFs that do not elicit a strong stress response could easily become subject to positive selection. Hence, our data support the notion that a pool of random sequences can easily provide the raw material for the *de novo* evolution of genes.

## 2. Materials and Methods

### Strains, plasmid and growth conditions

Three strains, E. coli K-12 DH10B (Neb 10-beta, NEB catalogue no. C3019H), E. coli B REL606 (Daegelen, Studier, Lenski, Cure, & Kim, 2009) and REL607 were used as backgrounds for this study. A multicopy expression vector, pFLAG-CTC™ (Sigma-Aldrich E8408), was used for the cloning and expression of candidate ORFs. Glycerol stocks were made by adding 700 µL of fully grown cultures into 700 µL of 50% glycerol and stored at −70ºC. Liquid media used for growth were: Lysogeny broth (LB) Lennox containing 10 g/L tryptone, 5 g/L yeast extract and 5 g/L NaCl, or Minimal medium (M9-Glucose) containing 33.9 g/L NaH2PO4, 15 g/L of KH2PO4, 2.5 g/L NaCl, 5 g of 1.8 M NH4Cl, 50 µL 1M CaCl2.6H2O, 1 mL 1 M MgSO4.7H2O and 10 mL 20% glucose. M9 media components were autoclaved and added separately to prevent precipitation and charring of glucose. The revival was done by streaking on agar plates with appropriate media (generally LB agar or M9 Glucose agar) supplied with 50 µg/mL Ampicillin (selection marker for plasmid) to obtain single isolated colonies that serve as clones for experimental replicates. Bacteria were generally incubated overnight for 16-18 h at 37ºC, shaking at a speed of 250 RPM (if shaking), unless otherwise mentioned.

### Cloning of selected candidates into E. coli strains

Selected candidate sequences were pulled out from the initial random library through PCR. Specific primers were designed for each sequence of interest (list provided in Supplementary Table 1) which were later used to amplify from the stored library plasmid DNA. Sequences were amplified using a 2step Phusion® high-fidelity PCR kit with the following PCR conditions: initial denaturation at 98°C for 30 seconds, followed by 30 cycles of – final denaturation for 98°C for 10 seconds with annealing at 72°C for 20 seconds. The final extension was performed at 72°C for 10 minutes, followed by indefinite cooling at 8-12°C. Phusion® PCR kit uses a high-fidelity Phusion polymerase with a 5’ to 3’ polymerase and 3’ to 5’ exonuclease activity. The amplified products were purified using a QIAquick PCR purification kit (Qiagen), then used for downstream cloning. The purified amplicons and purified vector DNA were digested using HindIII-HF™ and SalI-HF™ restriction enzymes for 1 hour at 37ºC, followed by purification using a QIAquick PCR purification kit. Purified products were ligated with 1 µL T4 DNA ligase (protocol as per NEB®) for 10 minutes at room temperature (benchtop) using a 3:1 insert to vector ratio. Ligation products were transformed in already competent cells background strains using chemical transformation. Commercial competent cells were used for the K-12 DH10B strain (NEB® 10-beta high efficiency). Transformation of B REL606 and B REL607 was achieved via the chemical competence method. For this, cells were prepared by growing cultures overnight in 4 mL LB medium and inoculating 500 ml of the pre-culture into fresh 200 mL LB medium and allowed to grow at 37°C, 250 rpm until an OD600 of 0.45-0.55 was reached. The culture was collected in four 50 mL FalconTM tubes and centrifuged for 10 m at 4°C, 3000 rpm. Pellets were gently resuspended in 1 mL chilled TBF-I solution (30 mM KOAc, 100 mM RbCl, 50 mM MnCl_2_ and 10 mM CaCl_2_) and filled up to 15 mL with the same. Tubes were incubated on ice for 1 hour followed by centrifugation for 10 m at 4°C, 3000 rpm. Pellets were then resuspended in 4 mL TBF-II solution (10 mM MOPS, 10 mM RbCl, 75 mM CaCl_2_ and 15% Glycerol).

Several transformation positive clones were freshly inoculated in 4 mL LB+ Amp media to prepare glycerol stocks the next day. The same colonies were also used for colony PCR to confirm the presence of insert. Colony PCR was done by taking a part of a fully-grown colony from the transformation positive plate and resuspending in 10 µL of sterile water. The suspension was heated at 98°C for 10 minutes and used as a template for subsequent 2-step Phusion PCR using specific primers. Amplicon confirmation was done using Sanger sequencing only if they showed a band corresponding to the insert (195 bp) on a 1.5 % agarose gel. Agarose gel electrophoresis was performed at 85-100 V for 30-45 minutes.

For confirmation of clones, pure amplicons were used for Sanger sequencing (Sanger, Nicklen, & Coulson, 1977). Common outer primers were used to amplify inserts (see supplementary Table 1). Output sequence files were analysed using Geneious prime (version 2019.1.3 or later) and CodonCode Aligner (version 7.0.1).

### Growth measurements

All growth curves were performed on Tecan M nano+ plate reader. Cultures from frozen stocks were streaked on appropriate media plates with ampicillin (50 µg/mL) and incubated overnight at 37°C (unless otherwise mentioned). The next day single colonies were inoculated in 200 µL of either LB or M9 medium with ampicillin in 96 welled plates and incubated with vigorous shaking for 16-18 hours. The following day, growth curves were set up by adding 2 µL of overnight culture to 200 µL of medium with ampicillin with or without 1 mM IPTG for induction of expression. Growth curves were usually performed for 15-24 hours with 5 minutes of orbital shaking before reading OD600 every 10 minutes at the desired temperature. The manual time series experiment was performed to obtain the colony-forming units in growing cultures. OD600 measurements in the plate readers measure turbidity and can be overestimated by factors like exopolysaccharide production. Strains were revived by standard procedures and growth measurements were started in larger volumes of 5 mL with a starting dilution of 1:100. After every hour of growth with or without IPTG induction at 250 rpm, cultures were plated on LA+Amp plates at an appropriate dilution and incubated overnight. Colonies were counted using a colony counter and CFU/mL at each time point was calculated. Growth curves were recorded for 13 hours.

### Population size estimation using manual CFU counts

Effects of NEG_Peps on the host viability were tested by manual growth estimation by CFU counts. Candidate strains were streaked on LA Ampicillin plates from respective glycerol stocks and incubated overnight at 37°C. The next day, 3 single colonies were inoculated in 4 mL LB Ampicillin and incubated at 37°C at 250 rpm overnight (16 hours). The following day, 40 µL from the overnight culture was inoculated into 4 mL LB Ampicillin tubes (1 in 100 dilution). Cells were allowed to grow for about 3-4 hours until the OD_600_ reached ∼0.4 (exponential phase). The uninduced time point (T0) was plated at an appropriate dilution to obtain a range of 30-300 colonies on LA ampicillin plates. Subsequently, the growing cultures were induced by adding 1mM IPTG (activates NEG_Pep expression) and cells were plated at appropriate dilutions after 5 min, 15 min, 30 min, 1 hour, 6 hours and 10 hours of induction. The observed colony counts were used to estimate CFU/ml at each of the above-mentioned time points.

### Competition experiments

Competitive fitness assays were performed using two E. coli strain backgrounds: B REL606 and B REL607 as described previously (Lenski et al., 1991). REL606 and REL607 are the ancestral strains of the *E. coli* long-term evolution experiment, which differ by a single point mutation in the arabinose utilisation (araA) gene of REL607, which allows it to metabolise arabinose. The two competitors can be distinguished by their arabinose utilisation phenotypes; (REL606) Araand (REL607) Ara+ that produce red and white colonies respectively on Tetrazolium agar (TA) indicator plates. The two backgrounds were transformed with our candidate ORFs on the pFLAG vector. First, strains were streaked on M9 Glucose Ampicillin agar plates and incubated at 37°C (unless otherwise mentioned) to obtain single colonies. Then, 4-8 colonies were inoculated the next day in 4 mL M9 Glucose Amp media per competitor and allowed to grow at 37°C for 18 hours, shaking at 250 rpm. On the subsequent day, a 1:1 volume of overnight cultures was used from each competitor strain and mixed thoroughly in a 96-welled plate (50 µL each strain). After mixing them in a 1:1 ratio, 40 µL of the mixture was inoculated in a fresh 4 mL M9 Glucose Amp media (with IPTG) and incubated for 24 hours at 37°C with shaking (250 rpm). The two competitor strains were allowed to grow for 24 hours at 37°C. Simultaneously, the mixture was plated (T0) on TA+Amp plates at an appropriate dilution such that the colony count was between 30-200 (statistical significance range) and incubated at 37ºC overnight. After 24 hours (T24), cultures were taken out and plated at an appropriate dilution and incubated at 37ºC overnight. For 48-hour competitions, 40 µL culture from the 24-hour tubes was transferred into fresh 4 mL media and allowed to grow for the next 24 hours at the same growth conditions. Colonies from the previous day were counted where a 50-50 ratio was expected. The next day, colonies from T24 were counted and the relative fitness of strains was determined.

Relative fitness was calculated as described in (Lenski, Rose et al. 1991):

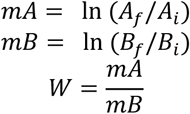

Where A and B are the two competing strains, i and f are the initial and final population densities (CFU/mL) of each competitor and W is the relative fitness of strain A w.r.t B. Ln is the natural logarithm. Malthusian parameter (m) of a strain (A or B) reflects population density changes over time.

### Total RNA extraction

Agar plates with appropriate media were streaked for different strain samples to be tested, were incubated at appropriate temperatures (37°C or 40°C) under static conditions. Different media (LA+Amp or M9+Amp) and temperatures were used but kept constant throughout the experiment. Similar culture and temperature conditions were used to acclimatize the strains to a particular environment in which they were being tested. After about 20-22 hours, single colonies (in 3-5 replicates) from the streaked plates were inoculated into 5 mL of liquid media (same as the one used in agar plates) followed by incubation at appropriate temperatures with shaking at 250 rpm overnight (16-18 hours). Inoculated tubes were allowed to grow until they reached the exponential phase, determined by measuring the optical density. An OD_600_ of 0.4-0.5 (standardized before the experiment) was considered adequate for the exponential phase of cultures. At this point, 500 µL of cultures were aliquoted, and 1 mM IPTG was added to the remaining cultures (4.5 mL). Aliquoted cultures were centrifuged for 5 minutes using a table-top mini centrifuge at maximum speed (13000 rpm) and pellets were stored at −20°C after treatment with RNAprotect reagent. The remaining cultures, induced by adding 1 mM IPTG, were immediately moved back to the shaker following the first aliquot and allowed to grow further. Another set of aliquots were taken after 1 hour of letting cultures grow under induction. Pellets were frozen in RNAprotect were used for RNA extraction within one week. Cultures aliquoted from different time points were vigorously (5 sec vortex) resuspended in 2 volumes equivalent of RNAprotect bacteria reagent (for 500 µL cultures, 1 mL reagent was added). The mixtures were incubated for 5 minutes at room temperature and then centrifuged for 10 minutes at 5000 x g. Supernatants were decanted and pellets were stored at −20°C for up to one week. RNAprotect bacteria reagent enters the cells and protects the RNA from denaturation. Total RNA extraction was performed using the RNeasy® Mini kit (Qiagen, Catalogue no. 74106) following the kit protocol. Final elution was performed with 40 µL of pure water. Total RNA samples were stored at −70°C until further use.

### Hybridization and feature extraction using E. coli microarray chips

RNA labelling and microarray hybridization were performed by following the supplier’s protocol of the chips (Agilent). The labelling kit generated cyanine labelled cRNAs which were amplified using the WT kit primer mix (mixture of oligo dT and random nucleotide-based T7 promoter primers), generating cRNA from samples. The provided spike-in controls were also labelled and amplified with the samples. Labelling, hybridization, washing and scanning were performed using the standard protocol from the

Agilent user guide (Low Input Quick Amp WT labelling kit). We performed one-color microarrays with the commercially available Agilent E. coli microarrays 8×15K, P/N G4813A, design ID 020097 with complete gene probes list. All experiments were run in triplicates.

### Differential expression analysis

We used the Limma software package on R to analyze the microarray data. All analyses are based on comparing expression level differences to RNA from the induced pFLAG empty expression vector. Top candidate genes were extracted from the respective expression tables produced by Limma analysis. We focused the Gene Ontology (GO) analysis on the genes with log2fold expression > 1 using Panther (version 16.0) (Mi, Muruganujan, & Thomas, 2013) or EcoCyc (Keseler et al., 2017), which usually provided the same results.

## 3. Results

Individual candidate clones in the present study were selected from the random sequence library described in (Neme et al., 2017). The expression vector design for this library is shown in suppl. file S1. In the previous study, the focus was on full-length inserts, i.e., 150 random nucleotides flanked by essential transcription and translation signals along with an N-terminal FLAG-tag. To allow tracing of the vector effect in the context of the other clones, we expanded the analysis to all ORFs, i.e., including the ones with a premature stop codon (Fajardo & Tautz, 2021). Candidate ORFs for this study were selected from this re-analysis to represent a range of log2fold changes and representations in the library. We chose six clones each with positive (POS) and negative (NEG) effects on cell growth. In addition, we chose five clones that had shown no significant (NS) growth change in the overall analysis of (Fajardo & Tautz, 2021). All clones code for full-length peptides except POS_Pep4b, 6 and 7, which have a premature stop codon (hence produce a shorter peptide). Figure 1A depicts these clones in the overall cloud of clones for the different growth cycle comparisons, Figure 1B show the individual trajectories of these clones. The sequences of all clones are provided in suppl. file S2. We find that the empty vector, which expresses a 38 amino-acid long polypeptide, shows only a small non-significant negative frequency change, i.e., at least in the bulk experiment, it does not have a strong negative effect on its own (Figure 1 pFLAG).

**Figure 1.**
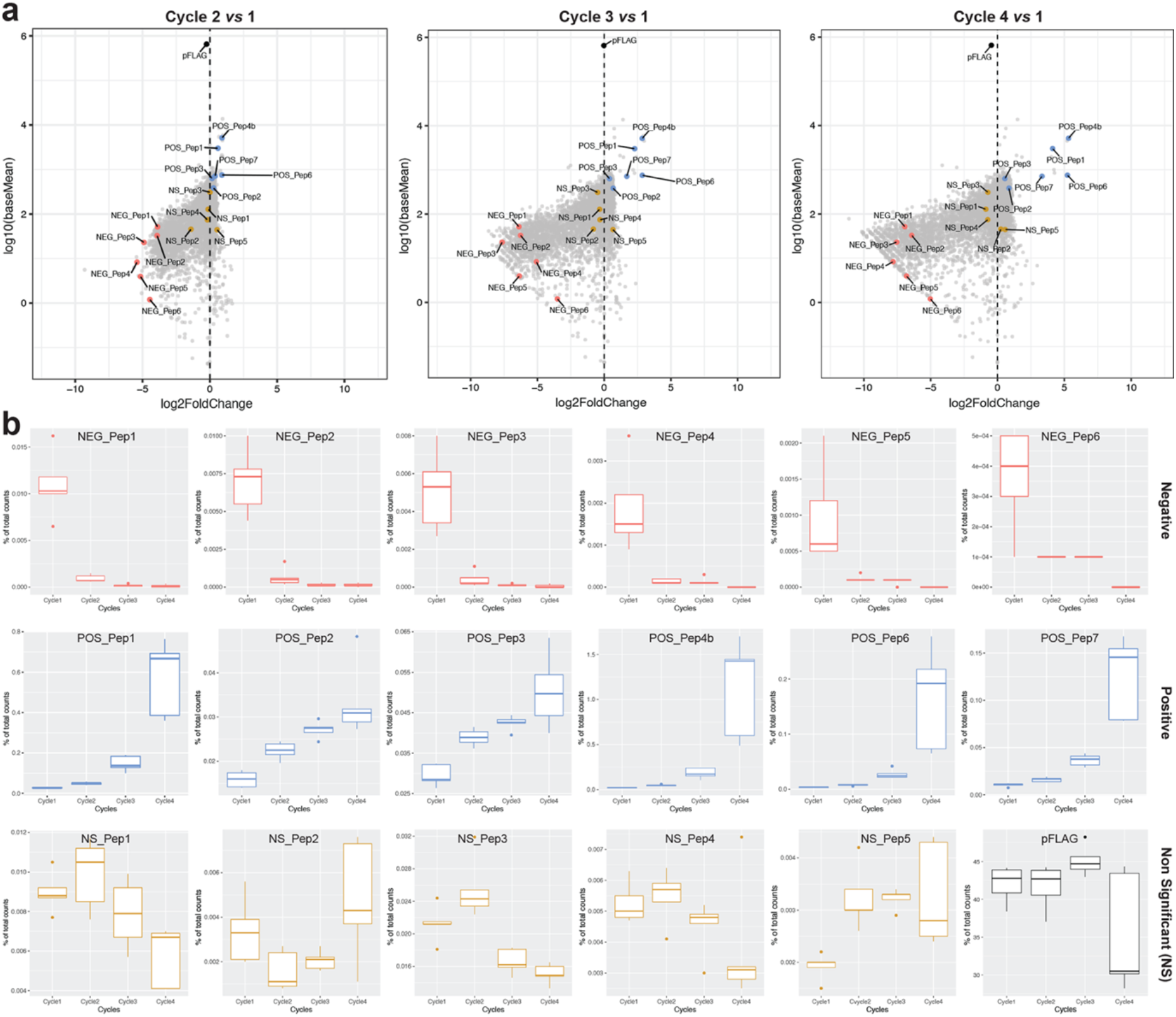
Performance characteristics of the candidate clones in the bulk experiment. Data are based on the deep sequencing experiment described in (Neme et al., 2017). **(a)**Frequency changes of candidate clones during the growth cycles of the random sequence library in the background of all analysed clones. Plots are based on the statistical analysis of clone frequency changes with DESeq2 (Love, Huber, & Anders, 2014), comparing mean counts of clones in the starting library (cycle 1) versus fold-changes at cycles 2, 3 and 4 (4 serial passages) for the deep sequencing experiment in (Neme et al., 2017) and its reanalysis in (Fajardo & Tautz, 2021). The Y-axis shows the average read counts across the replicates (log10(baseMean)); the X-axis is the relative change in clone frequency (log2FoldChange) at each cycle versus the counts at cycle 1. Candidates that are the focus of this study are highlighted with coloured dots. We categorise the candidates into three classes - NEG_Pep = negative (decreased in frequency - padj<0.05 and negative log2FoldChange value), NS_Pep = neutral (no significant change in frequency - padj>0.05), POS_Pep = positive (increased frequency - padj<0.05 and positive log2FoldChange values) and pFLAG = empty plasmid (black dot). Note that the overall effect of the empty plasmid in this experiment is within the range of the effects of the neutral peptides, but this can vary somewhat between experiments (Fajardo & Tautz, 2021). **(b)** Individual frequency trajectories of the candidate clones during the four cycles of growth. NEG clones show the expected ‘down’ pattern, POS clone the expected ‘up’ pattern, while the NS clones do not change much during the cycles. Boxplots show medians with lower and upper hinges corresponding to the 1^st^ and 3^rd^ quartiles.

### 3.1.1 Deleterious effects of negative clones in *E. coli*

We first focused on the growth characteristics of the negative clones. Growth curves were recorded for each clone under induced and non-induced conditions. All six clones show an initial growth delay after induction of the peptide before they resume growth (Figure 2A). The duration of the lag times differs slightly between the different clones, with NEG_Pep3 and NEG_Pep4 being particularly variable between replicates (Figure 2B). Growth resumes after the lag, but the rates remain significantly lower than the respective non-induced controls for five out of the six candidate clones (Figure 2C).

**Figure 2.**
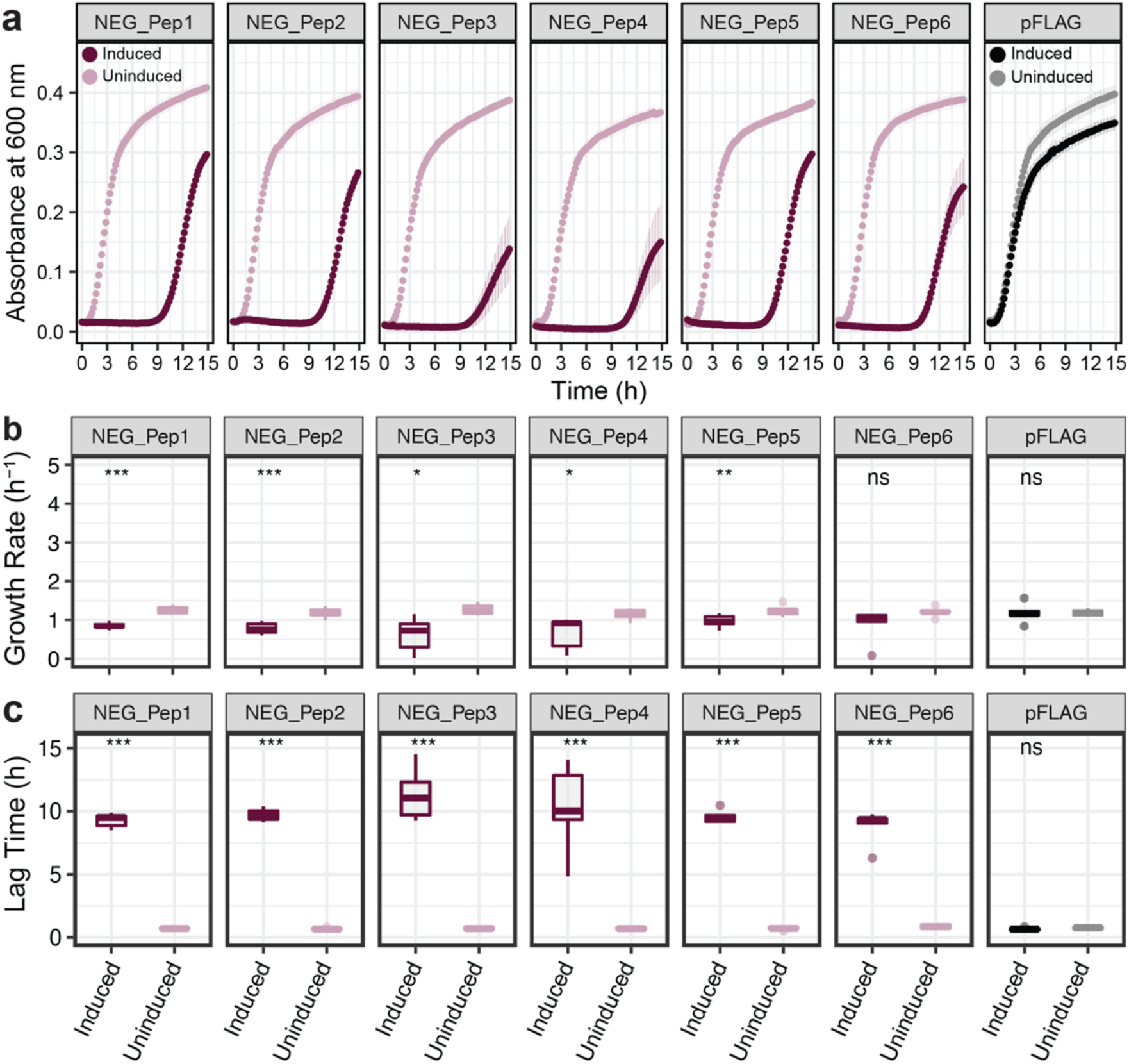
Growth rate comparisons for NEG peptide clones compared to empty vector. (**a**) Growth curves without IPTG induction are depicted as light colours and with IPTG induction are in dark colours. OD600 was recorded every 10 min with at least five replicates for each clone and condition. Dots represent means and whiskers, showing the standard error of the mean (SEM) across replicates. (**b**) Growth rate comparisons between the six clones after the end of the lag phase. (**c**) Lag time comparisons between the six NEG clones, measured as the time from the start of the experiment until the start of the exponential growth. Boxplots show medians with lower and upper hinges corresponding to the 1^st^ and 3^rd^ quartiles. Student’s t-test was performed with p-values as follows: *** = P < 0.001, ** = P < 0.01, * = P < 0.05, ns = P > 0.05.

We determined viable cell numbers during the lag phase to better understand the underlying cause of the growth delay. For this, we grew the bacteria until the log phase before adding IPTG. Samples were then taken at different times (up to 10 hours) and were plated on LB Ampicillin plates without IPTG (see methods). Figure 3 shows that viable cell numbers do not go down much in the first hour after IPTG induction, indicating that the cells are getting only growth-arrested but not immediately killed by the IPTG induction. However, viable cell numbers drop by two orders of magnitude by ten hours (Figure 3).

**Figure 3.**
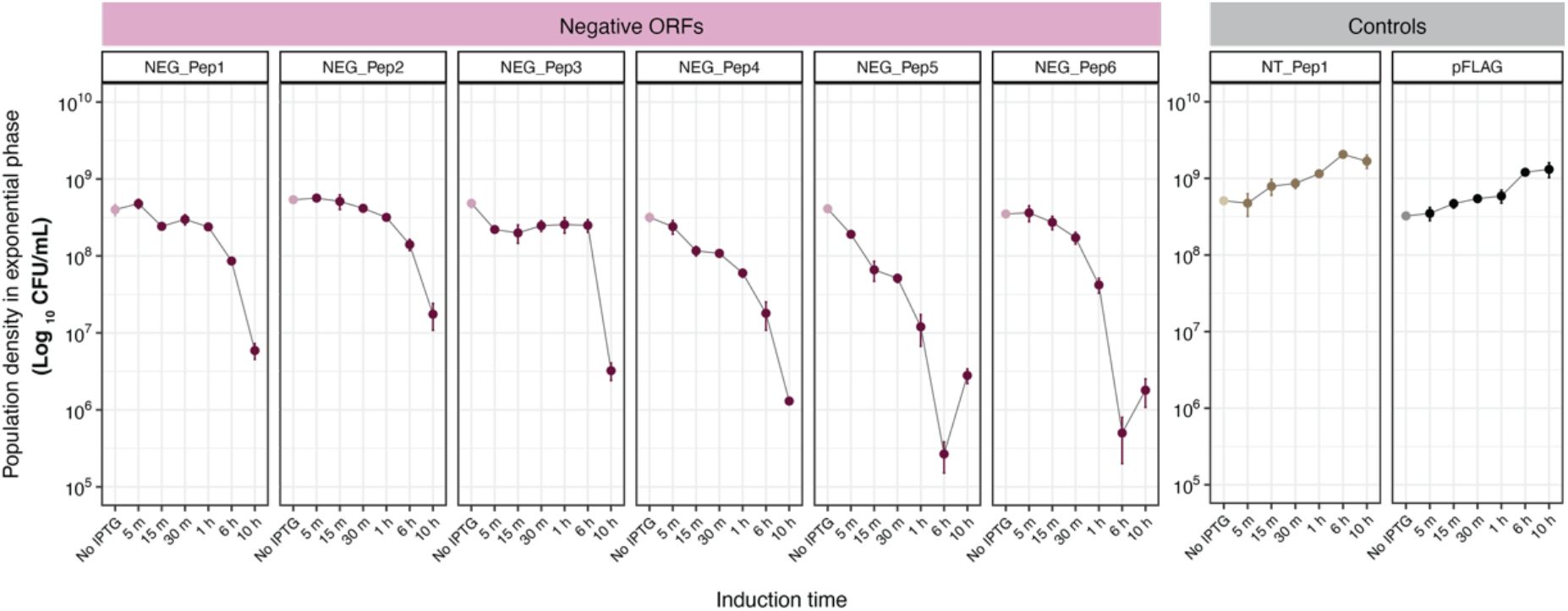
NEG peptide production causes a decline in the population of exponentially growing host bacteria. Colony counts (CFU/mL) of bacteria in the log phase, expressing either negative, neutral or pFLAG peptides at different times after IPTG addition, shows a decline in the NEG_Pep strains. Growth of the NT_Pep and pFLAG controls, on the other hand, is not affected by IPTG induction. Each panel’s light color dots represent growth under no induction, followed by IPTG addition (darker dots). Three replicates were done for each time point; whiskers in each dot represent SD.

To assess whether the negative effects of these clones are due to the expressed protein or mRNA, we tested six corresponding in-frame STOP codon versions of the clones that should express only the first three vector-encoded amino acids (suppl. file S2A). Western blots were used to show that the respective clones do not express the peptides anymore (data not shown). We found that four out of the six NEG clones still showed a growth delay due to the expressed RNA (suppl. File S3), implying that the RNA alone can negatively affect growth.

### 3.1.2 Transcriptomic response to NEG expression

Monitoring the *E. coli* RNA expression changes after IPTG induction is a way to study cellular responses. We used standardized *E. coli* microarrays to assess gene expression changes (Agilent). RNA was harvested 1h after induction (i.e., when most cells are arrested but viable) and expression changes were related to the RNA expression from cells carrying the induced vector plasmid without the insert.

The overall PCA indicates that each clone and their corresponding STOP codon versions show unique responses, whereby the main differences along PC1 correlate with peptide expression versus RNA expression (Figure 4a). Given a large number of genes with significant changes, especially for the clones expressing the peptide, we decided to focus on the more detailed analysis on the genes with the highest fold-change values (Figure 4b). Heatmap (based on log2 FC) of these genes shows an apparent clustering of NEG peptides versus the STOP codon versions of the clones (Figure 4c). This distinct picture suggests very different transcriptome reactions for expressing the proteins versus RNAs for these clones. But apart from these significant differences, it is also clear from the heatmap that there are distinct clone-specific differences.

**Figure 4.**
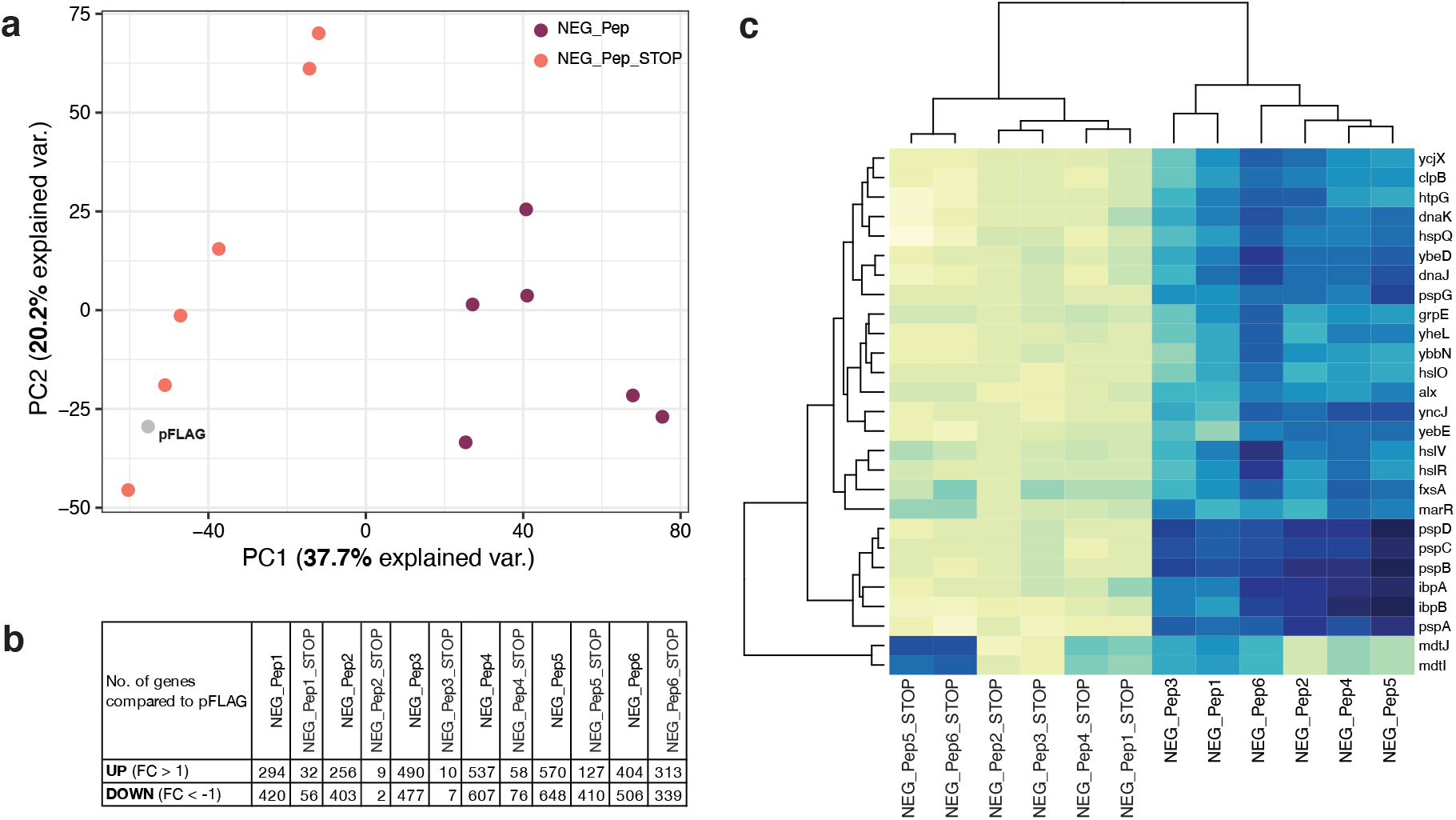
Transcriptomes of NEG peptides show differential up-regulation of genes compared to their stop codon versions. **(a)** Principal component analysis (PCA) of average expression from 3 replicates for each sample (as colored dots) are shown with PCs 1 and 2. **(b)** Number of genes above and below log2 fold change (FC) of 1 and −1 are shown here respectively. **(c)** Top highly expressed genes in all NEG_Peps are represented in the heatmap with dendrograms highlighting the differential expression in their corresponding stop codon versions. The X-axis shows the sample names and Y-axis lists the top genes (log2 FC > 4) of NEG_Peps from the expression analysis.

We used the gene lists of the top over-expressed genes to retrieve GO enrichment information from Panther GO (Ashburner et al., 2000; Gene Ontology, 2021). The top enriched categories overlap strongly for all six candidates (Figure 5). They include, in particular, stress response genes in various combinations, including protein folding. Only NEG_Pep3 shows fewer of these genes enriched. None of the stop codon versions of the peptides shows enrichment for these stress response genes; two show no enrichment for any GO category, one (NEG_Pep1_STOP) for “galactitol metabolic process” and the others for efflux transport processes (Figure 5). We conclude that a major consequence of the expression of the six NEG peptides is a strong stress response, with some modulation (e.g., in the case of NEG_Pep3). Furthermore, the stop codon versions of the respective candidate clones do not elicit the stress response. However, at least some of them cause almost equal growth inhibition as their corresponding peptide expressing clones, substantiating the notion that the RNA itself can cause effects on growth.

**Figure 5.**
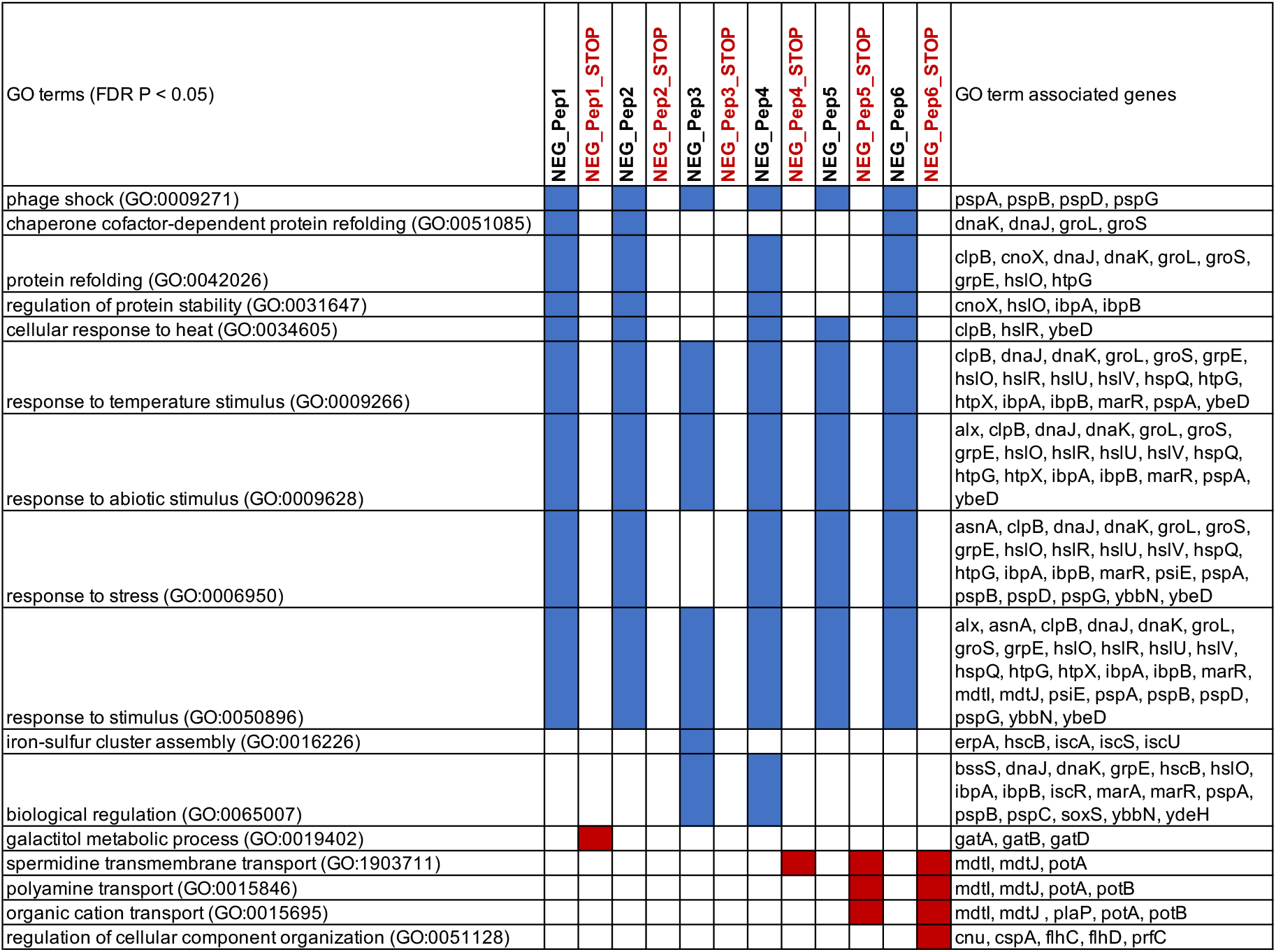
Major GO enrichment categories for the genes with the highest fold change for NEG peptides. Categories are listed with FDR p <0.05; redundant categories were removed. GO terms were extracted using the GO enrichment analysis tool using Panther version 16.0. Blue and red marks indicate that the respective GO categories were found for the NEG_Pep listed in the respective column. The GO term-associated gene list is cumulative for all NEG_Peps that show the respective GO term.

### 3.1.3 Growth characteristics of POS clones

In the second set of experiments, we focused on the further analysis of the positive clones. First, we assessed the growth characteristics of the positive clones in comparison to the NS candidates. Growth curves were recorded for each clone under induced versus non-induced conditions. Growth rates were determined during the exponential phase for each of them, but we did not find significant growth differences between the non-induced and induced conditions for any of them (suppl. file S4).

However, measurement of maximum growth rates of cultures during the exponential phase provides only a proxy for fitness. To increase the sensitivity of detection of growth differences, we performed competitive fitness assays in two strain backgrounds (*E. coli* B REL606 and *E. coli* B REL607) using the red-white selection as described in (Lenski, Rose, Simpson, & Tadler, 1991). The competitive fitness is calculated by estimating the population densities (CFU/mL) of each competitor at the beginning (T0) and the end (2 growth cycles) of the competition. Relative fitness and selection rates were calculated using Malthusian parameters of each competitor as described before (Lenski et al., 1991; Travisano, 1997). Genotypes with higher fitness produce more descendants and out-compete their less-fit competitors.

For the competitive fitness assays, each competitor was mixed in a one-to-one ratio from a fully-grown overnight culture (∼10^9^ cells/mL) and allowed to compete in IPTG supplemented minimal medium (M9+Glucose) for two 24-hour cycles. Note that competitive fitness assays measure the colony numbers as opposed to the growth rates. Swapped background strains were also engineered and tested to eliminate background related effects.

To avoid the problem of a possible negative effect caused by the peptide expressed by the empty vector (Knopp & Andersson, 2018), we used one of the NS candidates (NS_Pep3) in the comparative analysis of growth rates in the two strain backgrounds (see suppl. file S5 to support this choice). Most peptides identified as positive in the overall analysis also show a fitness advantage in the competition experiments (Figure 6). However, the effects are weak for POS_Pep1 and POS_Pep7, although both are very competitive in the bulk experiments (Figure 1). This suggests that the experimental conditions (minimal medium versus LB and two cycles versus four cycles) influence the performance of the clones. Overall, however, the experiment supports the notion that random peptide expression can lead to a growth advantage for the respective hosts.

**Figure 6.**
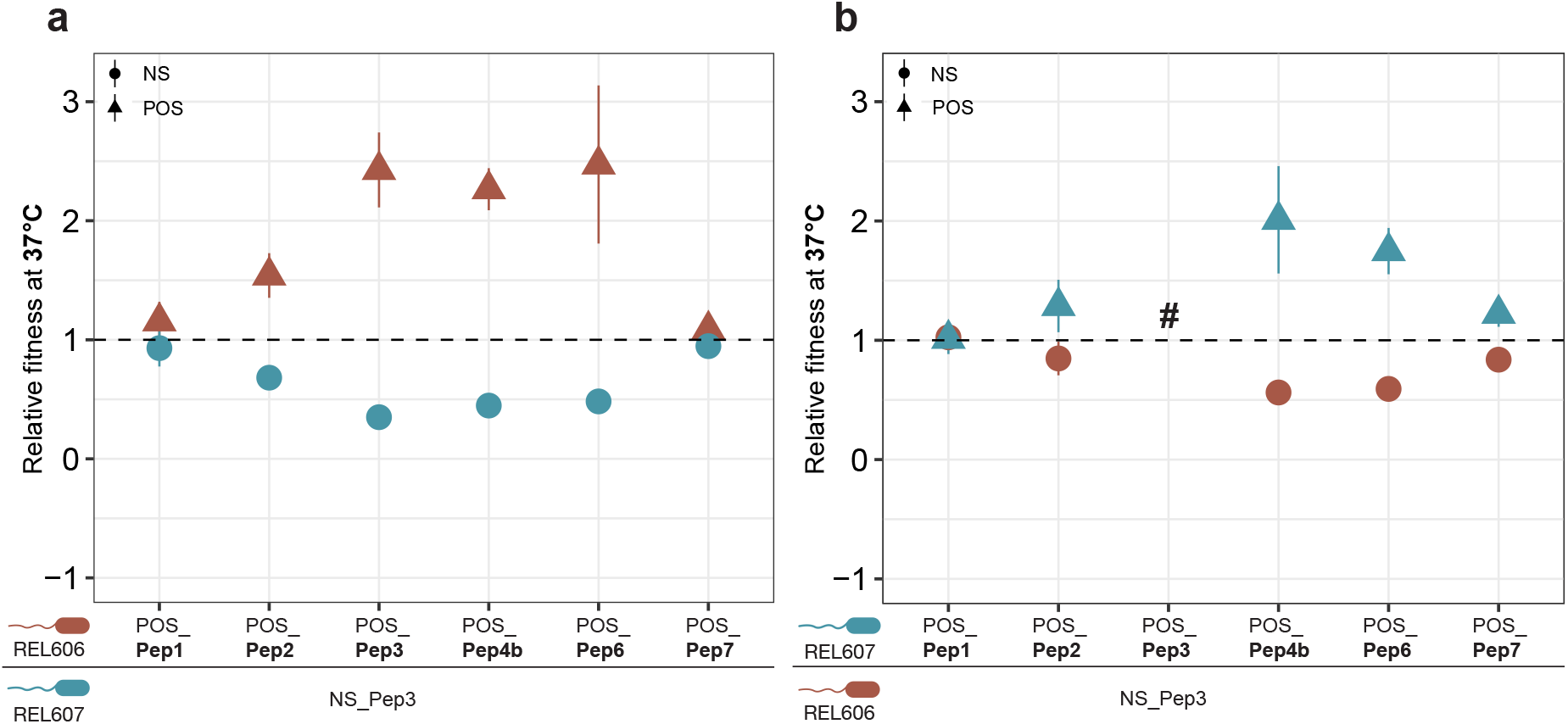
Competitive fitness analysis for six POS peptide clones. The respective plasmids were cloned into the *E. coli* B REL606 and REL607 ancestor backgrounds that produce red and pink colonies, respectively, on TA indicator plates (see Methods). The competitor in each experiment was the NS_Pep3 clone. (**a**) and (**b**) represent the experiments for swapped backgrounds to ensure that there are no background-specific effects. The symbol # represents missing data. At least four replicates were used for each experiment. Whiskers represent the SEM.

### 3.1.4. Transcriptomic response to POS expression

Using the above-described microarray system, we have also studied the transcriptomic responses of the POS peptides. These show a very different pattern compared to the NEG peptides. We generally observe a much weaker response at the transcriptional level and no activation of the general stress genes typical for the NEG peptides (see above). There is no apparent commonality between the clones; each shows different top GO categories for the relatively small sets of genes with log2 FC over-or underexpression (Table 1), including two without any GO enrichment (POS_Pep2 and POS_Pep7).

**Table 1.**
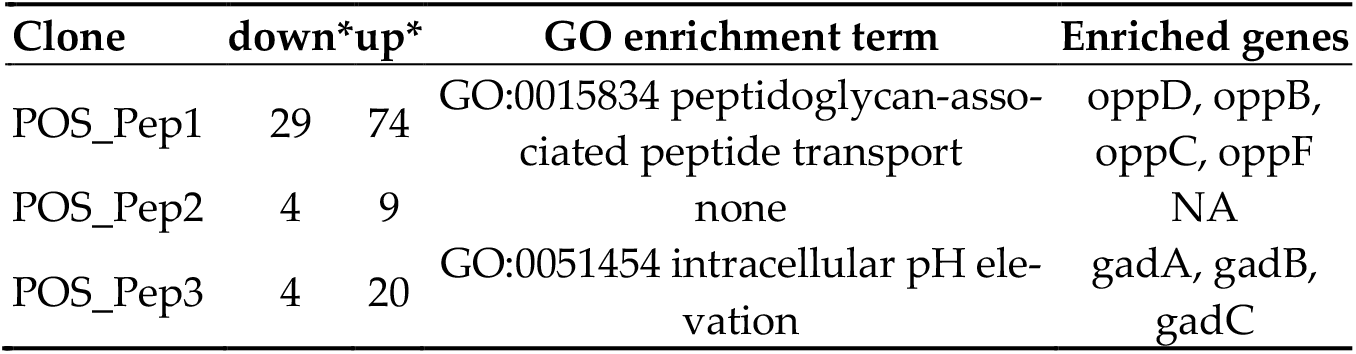
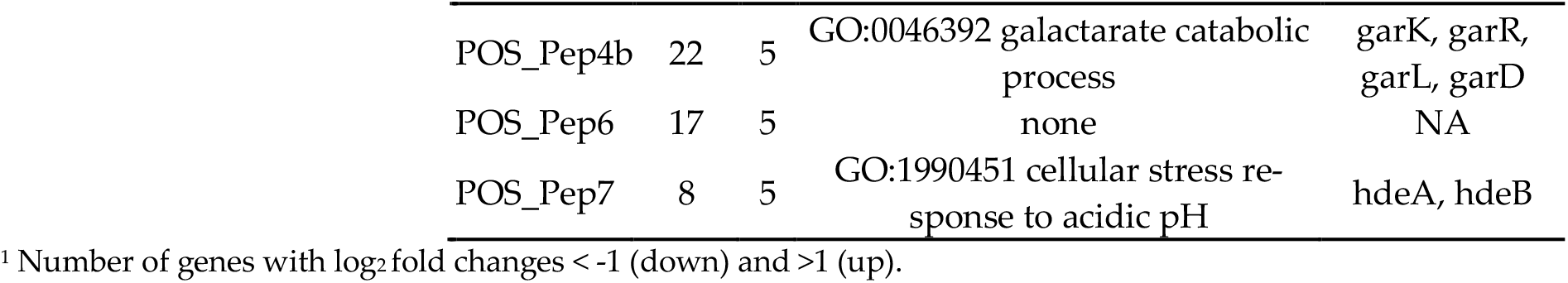
Transcriptomic responses of POS peptides.

## 4. Discussion

Studying the effects of expression of random sequence ORFs in cells can be considered a proxy towards understanding the evolution of new gene functions out of non-coding sequences (Neme et al., 2017; Weisman & Eddy, 2017). Here we have focused on a subset of candidate clones that broadly represent negative, neutral and positive categories when tested in the background of the growth of the whole library.

We find that the set of negative clones elicit a rather generic response in the cells, characterized by the strong upregulation of stress response genes, also known from clones that are used for recombinant protein production purposes (Gasser et al., 2008; Jurgen et al., 2000). Modulation of stress-related gene expression facilitates rapid adaptation in rapidly changing environments (Hengge-Aronis, 2002). This leads to an arrest in cell growth and eventually the death of a large number of cells in a population. Interestingly, a fraction recovers from this arrest after a certain lag, which is seen as a jump in growth rates back to normal. While we did not further explore how this switch back to normal growth is achieved, it can explain why the clones are not entirely lost in the bulk experiments with the whole library. Although the activation of the stress response is a common theme among these clones, they also have more specific effects partly. NEG_Pep3 and NEG_Pep4, for example, show additional sets of differentially regulated genes in response to IPTG induction.

Our experiments with the in-frame STOP codon version of the NEG clones provide insights into the effect of RNA versus protein in these random sequence clones. For two clones, we find a complete recovery of the expected growth; for another two clones, we see only a partial recovery, and the final two clones still show a similar lag as their corresponding peptide expressing versions (suppl. file S3). Interestingly, these latter two clones (Neg_Pep5_STOP and NEG_Pep6_STOP) do not elicit a stress response (Figure 4C), implying that another pathway mediates their effect on growth delay. This observation also indicates that it is not the overexpression that inhibits the host growth per se but that there are true RNA and protein-specific effects. The fact that growth delay can also be caused by RNA transcripts alone, i.e. independent of translation, was also found for clones used for recombinant protein production (Li & Rinas, 2021) and in experiments expressing different GFP RNA variants (Mittal, Brindle, Stephen, Plotkin, & Kudla, 2018). On the other hand, the re-analysis of (Fajardo & Tautz, 2021) has suggested that also the first three amino acids coded in these STOP codon versions of clones can have detrimental effects on their own, but depending on the rest of the RNA.

The positive clones show a distinct response. They do not affect the cell growth rates but show a competitive growth advantage when tested individually against a neutral candidate clone. The transcriptomic response of positive clones is much lower compared to the transcriptomic response of negative ones. Only a few genes are enhanced or repressed in positive clones, and there is no common theme among the GO categories for the enriched genes. In independent repetition experiments with slightly changed conditions, we have noted that even these responses were not stable and that different sets of genes came up in the top lists for the positive clones (data not shown). Hence, we conclude that the effects of the POS peptides are unlikely to be directly at the transcriptional level. The changes that we observe in the transcriptome may be secondary effects of interactions of these peptides within the cytoplasm.

Our study suggests thus that random sequence clones potentially interact with the cellular machinery of bacteria in a promiscuous manner. Although negative peptide clones have robust deleterious effects on the host, the clone-specific variation in gene expression may eventually provide insights into the individual effects of peptides. Note, however, that we have not tested the STOP-codon versions of these clones in the competition experiments, hence it remains at this point open whether the positive effect is due to protein or RNA expression or both. previous re-analysis of such positive clones has suggested that both RNA and protein can cause the effect (Neme et al., 2017).

## 5. Conclusions

Our analysis confirms the overall notion that a substantial fraction of random sequence ORFs does not have detrimental effects on cells, even when highly over-expressed. Moreover, at least a subset can even provide growth advantages to the cells in direct competition experiments. This proves their general potential for the raw material for new evolutionary adaptations, which always start with a growth advantage compared to other cells under given conditions. For example, Knopp and colleagues (Knopp et al., 2019) have studied whether random sequence peptides can convey antibiotic resistance and found several clones with such an effect. Functional analysis showed that the effect is caused via membrane depolarization, which decreases aminoglycoside uptake and thus provides a growth advantage under the presence of an antibiotic. However, when these clones were competed against the empty vector plasmid, they showed a growth disadvantage (Knopp et al., 2019), i.e., would be categorized as negative in our comparisons. Hence, any growth advantages or disadvantages triggered by random sequence clones need to be seen in the context of the competitors in the experiment (Weisman & Eddy, 2017). It would seem likely that growth differences measured in the bulk experiments with many competing clones would not be directly comparable to growth differences when only two specific clones compete. But given that we see for at least four of the POS clones derived from the statistical analysis of the bulk experiments also a clear growth advantage in pairwise competitions, these effects cannot simply be explained by compensation of expression vector effects. Interestingly, while beneficial point mutations are relatively rare in long-term evolution experiments in *E. coli* (Barrick & Lenski, 2013), the fraction of reproducible positive growth effects from random sequence clones is relatively high (at least around 10% of all clones), implying that the expression of a new peptide can more easily lead to an evolutionary innovation than point mutations.

## Author Contributions

Conceptualization, D.B. and D.T.; investigation, D.B.; data curation and analysis, D.B. and D.T.; writing—original draft preparation, D.B.; writing—review and editing, D.T.; supervision, D.T. All authors have read and agreed to the published version of the manuscript.

## Acknowledgements

We thank Dr Jenna Gallie for suggestions on experimental procedures and comments, Elke Blohm-Sievers for help with the microarray experiments and Johana Fajardo for sharing her re-analysis data. This research was funded through an ERC advanced grant to D.T. (NewGenes—322564) and institutional funds of the MPG to D.T. D.B. was a member of the IMPRS for Evolutionary Biology.

