## Supplementary figures and tables for "Effects of the expression of random sequence clones on growth and transcriptome regulation in *Escherichia coli*"

### Supplementary files

**suppl file S1**

Expression vector and insert design. The IPTG inducible expression vector pFLAG-CTC was used to clone random sequence ORF inserts into the multiple cloning site (MCS) between the HindIII and SalI sites, as shown on top. The scheme of the full-length peptide product expressed upon induction is shown at the bottom.

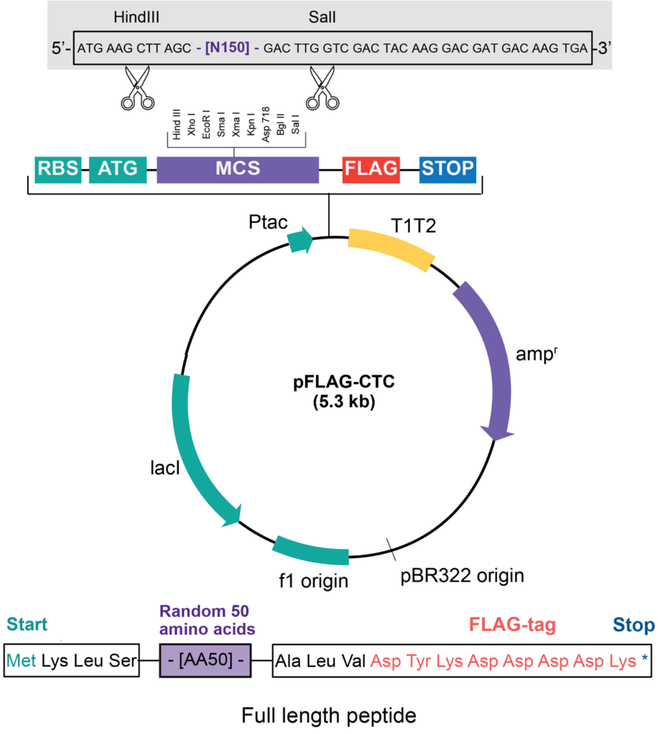


**suppl file S2**

Sequences of clones in this study: coding parts of the inserts are shown with their translation products. Constant parts derived from the vector sequences are shaded in grey.

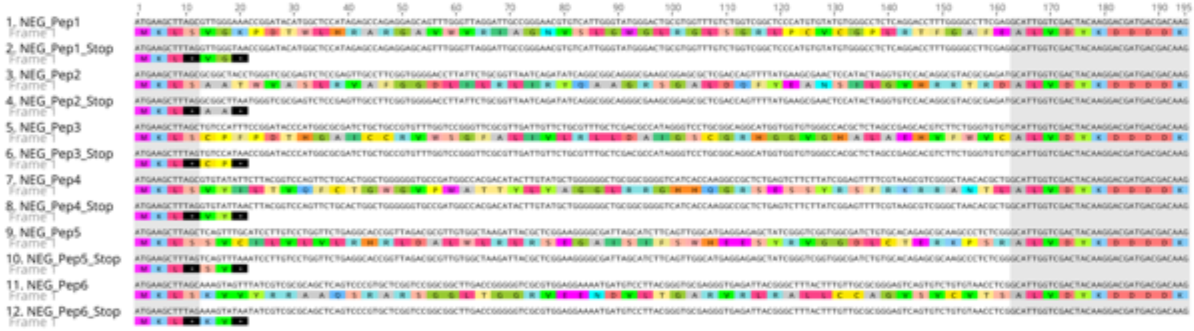

*S2A: NEG clones and their STOP codon variants.*

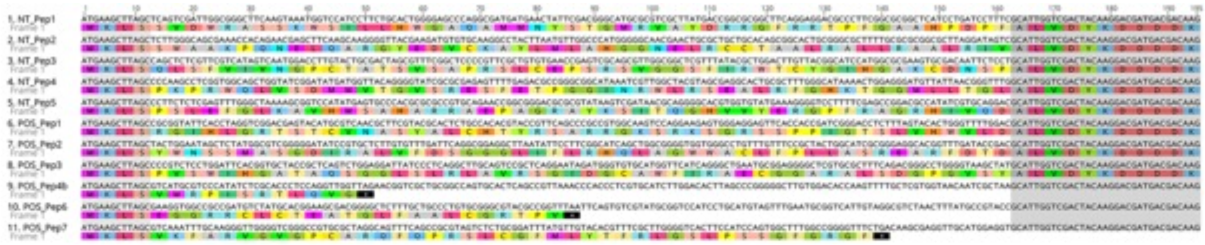

*S2B: NT and POS clones.*

### suppl file S3

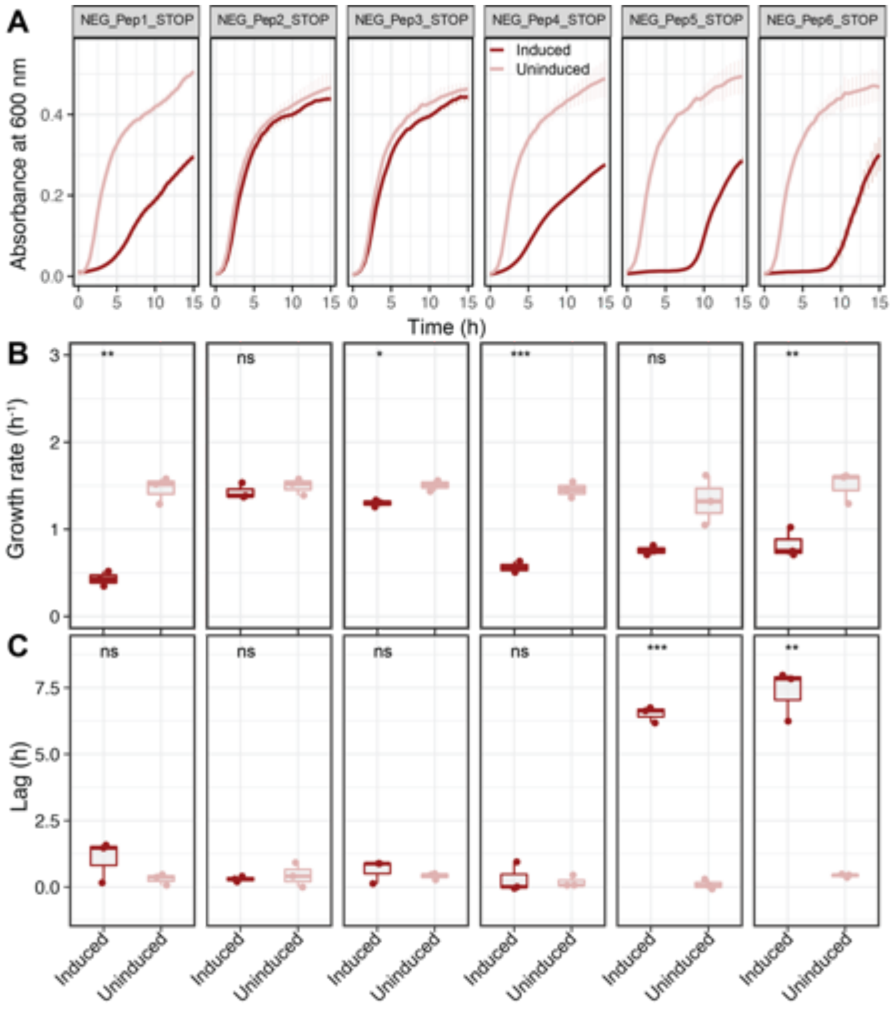


**Growth rate comparisons for NEG peptide STOP clones.** A) Growth curves without IPTG induction are depicted as light colours and with IPTG induction are in dark colours. OD600 was recorded every 10 min with at least five replicates for each clone and condition. Dots represent means and whiskers, which show the standard error of the mean (SEM) across replicates. B) Growth rate comparisons between the six clones after the lag. C) Lag time comparisons between the six NEG STOP clones, measured as the time from the start of the experiment until the start of the exponential growth. Student's t-test was performed with p-values as follows: *** = $P < 0.001$, ** = $P < 0.01$, * = $P < 0.05$, ns = $P > 0.05$.

**suppl file S4**

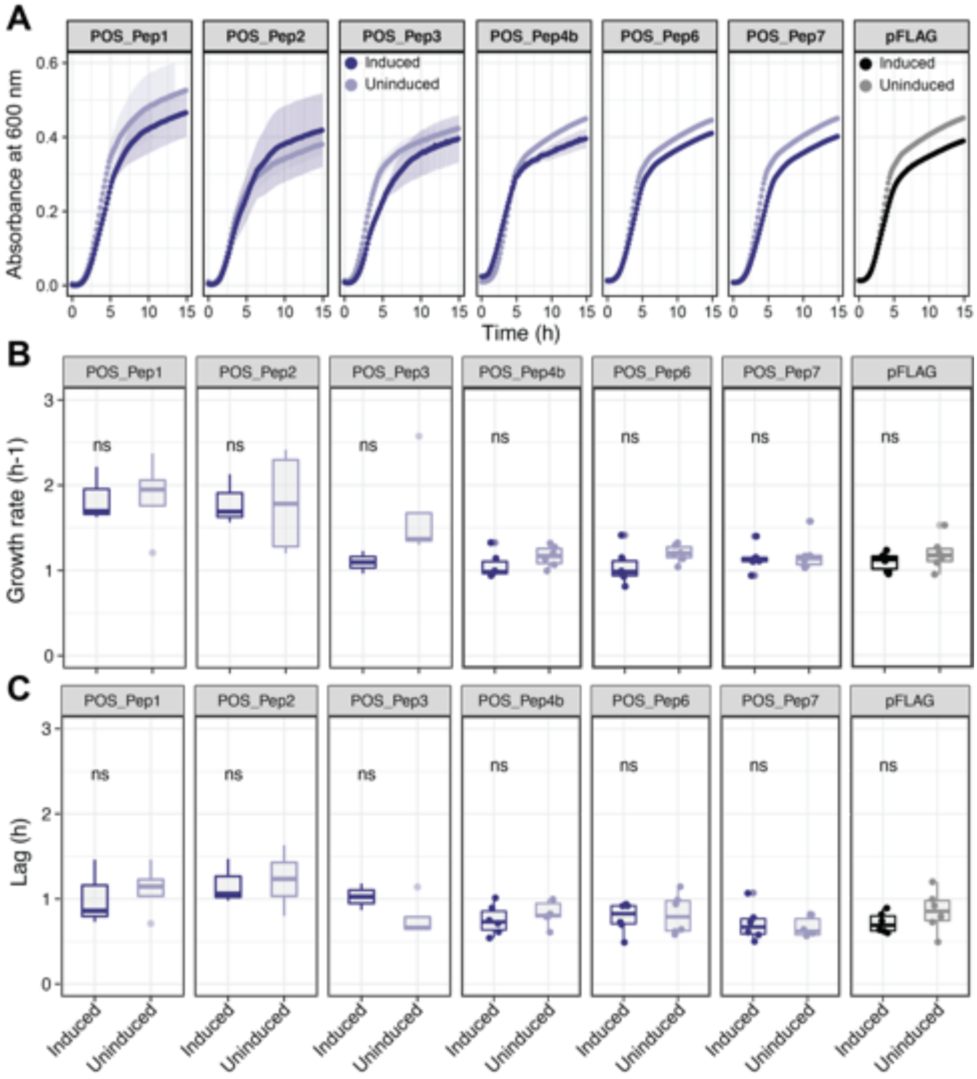


**Growth rate comparisons for POS peptide clones compared to empty vector.** A) Growth curves without IPTG induction are depicted as light colours and with IPTG induction are in dark colours. OD600 was recorded every 10 min with at least five replicates for each clone and condition. Dots represent means and whiskers, which show the standard error of the mean (SEM) across replicates. B) Growth rate comparisons between the six clones. C) Lag time comparisons between the six clones, measured as the time from the start of the experiment until the start of the exponential growth. Student's t-test was performed with p-values as follows: *** = $P < 0.001$, ** = $P < 0.01$, * = $P < 0.05$, ns = $P > 0.05$.

**suppl file S5**

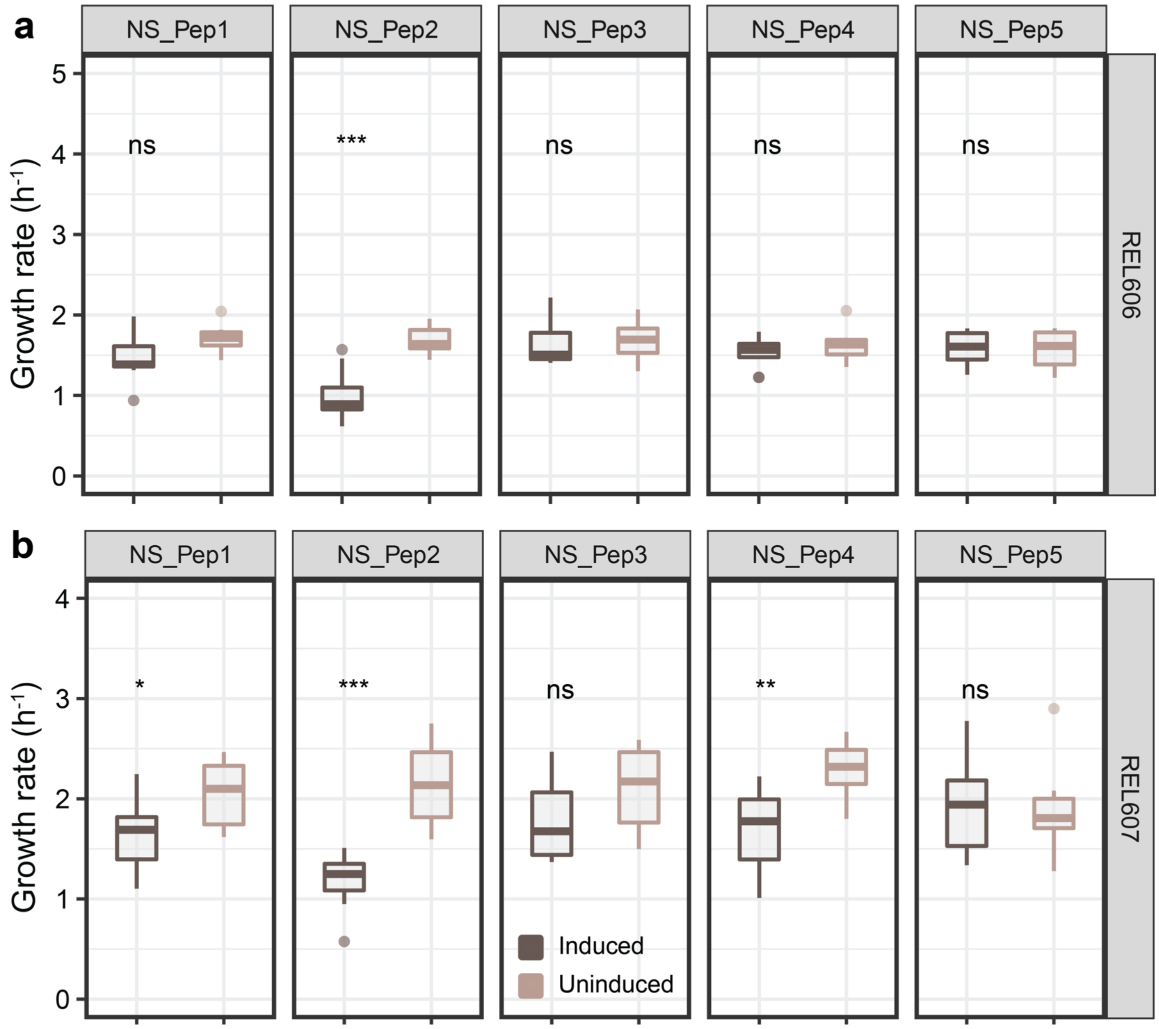


**Growth rate comparisons for NS peptide clones in two different backgrounds.** A) Growth rate comparisons between the five clones in REL606. B) Growth rate comparisons between the five clones in REL606.Student's t-test was performed with p-values as follows: *** = $P < 0.001$, ** = $P < 0.01$, * = $P < 0.05$, ns = $P > 0.05$.

**Supplementary Table 1**

Primer sequences used in this study

| Name | Primer | Sequence |
|---|---|---|
| NEG_Pep1 | Forward | ATGAAGCTTAGCGTTGGGAAACCGGATACATGGC |
| NEG_Pep1 | Reverse | GTAGTCGACCAATGCCTCGAAGGCCCCAAAGGTC |
| NEG_Pep1_STOP | Forward | ATGAAGCTTTAGGTTGGGTAACCGG |
| NEG_Pep1_STOP | Reverse | GTAGTCGACCAATGCCTCGAAGG |
| NEG_Pep2 | Forward | ATGAAGCTTAGCGCGGCTACCTGGGTCGCGAGTC |
| NEG_Pep2 | Reverse | GTAGTCGACCAATGCATCTCGCGTACGCCTGTGG |
| NEG_Pep2_STOP | Forward | ATGAAGCTTTAGGCGGCTTAATGGGT |
| NEG_Pep2_STOP | Reverse | GTAGTCGACCAATGCATCTCGCGTA |

| | | |
|---|---|---|
| NEG_Pep3 | Forward | ATGAAGCTTAGCTGTCCATTTCCGGATACCCATG |
| NEG_Pep3 | Reverse | GTAGTCGACCAATGCACACACCCAGAAGACGTGC |
| NEG_Pep3_STOP | Forward | ATGAAGCTTTAGTGTCCATAACCGGATACCC |
| NEG_Pep3_STOP | Reverse | GTAGTCGACCAATGCACACACCC |
| NEG_Pep4 | Forward | ATGAAGCTTAGCGTGTATATTCTTACGGTCCAGT |
| NEG_Pep4 | Reverse | GTAGTCGACCAATGCCAGCGTGTTAGCCCGACGC |
| NEG_Pep4_STOP | Forward | ATGGAAGCTTTAGGTGTATTAACTTACGGTCCAGT |
| NEG_Pep4_STOP | Reverse | GTAGTCGACCAATGCCAGCGTG |
| NEG_Pep5 | Forward | ATGAAGCTTAGCTCAGTTTGCATCCTTGTCCTGG |
| NEG_Pep5 | Reverse | GTAGTCGACCAATGCCCGAGAGGGCTTGCGCTCT |
| NEG_Pep5_STOP | Forward | ATGAAGCTTTAGTCAGTTTAAATCCTTGTCCTGG |
| NEG_Pep5_STOP | Reverse | GTAGTCGACCAATGCCCGAGAGG |
| NEG_Pep6 | Forward | ATGAAGCTTAGCAAAGTAGTTTATCGTCGCGCAG |
| NEG_Pep6 | Reverse | GTAGTCGACCAATGCCGAGGTTACACAGACACTG |
| NEG_Pep6_STOP | Forward | ATGAAGCTTTAGAAAGTATAATATCGTCGCGC |
| NEG_Pep6_STOP | Reverse | GTAGTCGACCAATGCCGAGGTTACA |
| POS_Pep1 | Forward | ATGAAGCTTAGCCGCGGTATTCACCTAGGTCGGA |
| POS_Pep1 | Reverse | GTAGTCGACCAATGCGTCCAAAACCCAGTGT |
| POS_Pep2 | Forward | ATGAAGCTTAGCTACTGGAATAGCTCTATGGCGT |
| POS_Pep2 | Reverse | GTAGTCGACCAATGCGTCGGTATCAAACCGT |
| POS_Pep3 | Forward | ATGAAGCTTAGCCCCGTCTCCTGGATTCACGGTG |
| POS_Pep3 | Reverse | GTAGTCGACCAATGCATAGCTTACCCCAGGC |
| POS_Pep4b | Forward | ATGAAGCTTAGCGTCATGCGTCCCATATCTCG |
| POS_Pep4b | Reverse | TCATCGTCCTTGTAGTCGACCAATGCTTAGCG |
| POS_Pep6 | Forward | ATGAAGCTTAGCGAAGGTGGCCGCCGAT |
| POS_Pep6 | Reverse | CATCGTCCTTGTAGTCGACCAATGCGGTACG |
| POS_Pep7 | Forward | ATGAAGCTTAGCGTCAAATTTGCAAGGGTTGGGGTC |
| POS_Pep7 | Reverse | TCATCGTCCTTGTAGTCGACCAATGCACCTC |
| Outer_pFLAG | Forward | GCATAATTCGTGTCGCTCAA |
| Outer_pFLAG | Reverse | AAAAGGGAATAAGGGCGACA |
